# A comprehensive bulk and single-cell transcriptional atlas of pediatric leukemias

**DOI:** 10.64898/2025.12.23.696262

**Authors:** Irina Pushel, Tarun KK Mamidi, Byunggil Yoo, Lisa A Lansdon, Daniel Louiselle, Margaret Gibson, Erin Guest, Nicole M Wood, Keith J August, Terrie G Flatt, Alan S Gamis, Tomi Pastinen, Midhat S Farooqi

## Abstract

Advancements in understanding the molecular factors driving pediatric leukemias have led to an ever-increasing volume and diversity of data being generated. Recent studies are moving beyond DNA-based profiling to incorporate transcriptional data, enhancing the characterization of these cancers and informing clinical decisions. However, many existing datasets focus on specific leukemia subtypes, limiting the extraction of broader clinically relevant insights. Here, we present a comprehensive dataset of bulk and single-cell transcriptional data from 69 pediatric leukemia patients, encompassing eight leukemias. This is one of the most diverse pediatric leukemia datasets published to date. Through comparative analysis and explainable machine learning, we demonstrate the utility of these datasets in improving pediatric leukemia characterization and supporting the integration of transcriptional data into clinical testing.

## Introduction

Recent advances in sequencing technologies have increased the use of sequencing data in cancer research and in clinical settings. The development of clinical assays using panel, whole exome, and whole genome sequencing has led to the identification of DNA variants driving cancers. Combined with existing clinical metrics, genomic data help identify cancers requiring more aggressive treatment as well as those which may respond to targeted therapies (J. Li et al., 2021; Tasian et al., 2015).

Despite these improvements, approximately 10% of leukemia cases remain with no clear genomic driver after DNA-based testing. This is particularly true of pediatric leukemias, which are rarer than their adult counterparts and often behave differently (Ahmed et al., 2018; Brady et al., 2022; Milan et al., 2019; Vogelstein et al., 2013). Some leukemias without a unifying driver mutation, such as Philadelphia-like acute lymphoblastic leukemia (Ph-like ALL), nonetheless show a unified set of molecular characteristics and transcriptional similarity to *BCR::ABL1* or Philadelphia-positive (Ph+) ALL. These cases are characterized by *CRLF2* and *ABL1/2* deregulation and are usually responsive to tyrosine kinase inhibitors (Reshmi et al., 2017; Roberts et al., 2014; Tran & Loh, 2016). This example directly demonstrates the value added by incorporating transcriptional data, in addition to genomic variant data, in the diagnosis and treatment of pediatric leukemias.

Increasingly, RNA-based assays are being incorporated into clinical practice for pediatric leukemia patients, through RNA-based fusion detection and gene expression classifiers (Hu et al., 2023; Vaske et al., 2019). To improve the utility of transcriptomic data in the clinic, several efforts have been undertaken to sequence and share pediatric cancer transcriptomes (Sanders et al., 2019; Vaske et al., 2019), including leukemias specifically (Gu et al., 2019; Iacobucci et al., 2025; Mumme et al., 2022; Pölönen et al., 2024; Tan et al., 2023). However, many of these datasets are limited to individual leukemias or small subsets and share only bulk or single-cell RNAseq data, but not both.

Here, we have generated bulk and single-cell transcriptional data from 69 patients that span different types of leukemias, including acute myeloid leukemia, B-cell acute lymphoblastic leukemia, and T-cell acute lymphoblastic leukemia. In addition to contributing data to a relatively sparse ecosystem, this dataset is one of the few to contain both bulk and single-cell RNAseq data for the same patients, allowing researchers to make direct comparisons between the two datasets and extrapolate findings to other datasets. The raw and processed data are accessible through the Childhood Cancer Data Initiative (CCDI) and single-cell data are additionally available in interactive form at the Single Cell Portal at https://singlecell.broadinstitute.org/single_cell/study/SCP3385/single-cell-profiling-of-pediatric-leukemia-patient-samples. Furthermore, we have used explainable machine learning to develop one of the first gene-expression classifiers for single-cell gene expression profiles of pediatric B-ALL, which is available at https://github.com/ChildrensMercyResearchInstitute/cm-scbll.

## Materials and Methods

### Sample sources and characteristics

This study utilized peripheral blood and/or bone marrow samples obtained from the Children’s Mercy Tumor Bank Biorepository (CRIB). Samples were collected at diagnosis, remission, and/or relapse, then cryopreserved until processing. Patient demographics and clinical characteristics are summarized in Table 1. Molecular subtypes were determined based on clinical genomic testing, including FISH, cytogenetics, and targeted mutation profiling using the CCDI submission guidelines. Some cases were later reclassified based on findings from the subsequent tests. Full patient demographic and clinical data available at https://datacatalog.ccdi.cancer.gov/dataset/CCDI-phs002529. This research was conducted in accordance with the Declaration of Helsinki and approved by the Children’s Mercy Institutional Review Board. All patient samples were collected with written informed consent of the parents/guardians and assent of patients.

**Table 1.** Performance metrics for ScPCA test and CM datasets. cells – cell level predictions; patients – patient level predictions.

| Dataset | Accuracy | Precision | Recall | F1 score |
| --- | --- | --- | --- | --- |
| Iacobucci et al. test data (cells) | 0.99 | 0.99 | 0.99 | 0.99 |
| CM samples (cells) | 0.75 | 0.88 | 0.75 | 0.81 |
| CM samples (patients) | 0.8 | 0.91 | 0.8 | 0.84 |

### Library preparation and sequencing

#### Bulk RNA Sequencing

Bulk RNA-seq libraries were prepared using the Illumina Stranded Total RNA with Ribo-Zero Gold library preparation kit. Paired-end sequencing was performed on an Illumina NovaSeq 6000 platform.

#### Single-Cell RNA Sequencing

Single-cell RNA-seq data were generated in two distinct batches—referred to as the pilot and s2022 batches. (i) Pilot batch: A total of 29 samples were multiplexed across three pools. Two captures per pool, each targeting 16,000-24,000 cells, were performed using a 10x Genomics Chromium Controller. Libraries were prepared using 10x Single Cell 3’ v2 chemistry and sequenced as pooled libraries on an Illumina NovaSeq 6000. (ii) s2022 batch: A total of 43 samples were multiplexed across four pools. Five captures per pool, each targeting 10,000 transposed nuclei, were performed using a 10x Chromium Controller. Libraries were prepared using the 10x Multiome ATAC + Gene Expression v1 chemistry and sequenced as pooled libraries on an Illumina NovaSeq 6000.

### Bulk RNAseq data analysis

Reads were aligned to the GRCh38 reference genome using STAR v2.6.0 (Dobin et al., 2012), and gene-level quantification was performed using kallisto v0.46.2 (Bray et al., 2016). Raw and processed RNAseq data, along with matched whole-genome and/or whole-exome sequencing data, are available via the Childhood Cancer Data Initiative at https://datacatalog.ccdi.cancer.gov/dataset/CCDI-phs002529. Downstream analysis was conducted in R v4.0.3 using DESeq2 v1.36.0 (Love et al., 2014), with sequencing batch included as covariate. For gene expression visualization, log-normalized counts were batch-corrected using the limma package v3.52.4 (Ritchie et al., 2015) to remove the ‘batch’ covariate. Differential gene expression analysis was performed using the likelihood ratio test (LRT), and genes with adjusted p-value < 0.05 were considered statistically significantly differentially expressed.

### Single-cell RNAseq data analysis

Raw sequencing data from the pilot batch were processed using 10x Genomics Cell Ranger (Zheng et al., 2017) v3.0.2, while the s2022 batch was processed using 10x Genomics Cell Ranger ARC 2.0.0, both aligned to GRCh38. For demultiplexing purposes, bcftools (Danecek et al., 2021) v1.10.2 was used to merge vcf files from whole genome or whole exome sequencing for all of the patients in a particular pool, generating a pool-specific multi-sample vcf file. This multi-sample vcf file was then used as an input to popscle demuxlet (Kang et al., 2017) commit 3e4a38b using default settings to demultiplex the data and assign each cell called by Cell Ranger to a patient sample. For s2022 data, demuxlet was applied independently to RNA and ATAC data for each capture.

Downstream processing was performed in R v4.0.3 using Seurat v4.0.2 (Hao et al., 2021; Satija et al., 2015). Cells were filtered to retain only those with (i) a demuxlet ‘singlet’ call based on RNA data, (ii) between 850 and 25,000 molecules per cell (nCount_RNA), and (iii) less than 20% mitochondrial reads. This filtering yielded a total of 144,747 high-confidence cells. Each batch was independently filtered and normalized using SCTransform in Seurat, then merged into a unified Seurat object for downstream analysis. Cell-type annotations were assigned using the SingleR v1.4.0 package (Aran et al., 2019) with the celldex Blueprint ENCODE reference dataset to assign most similar cell types for each cell in the dataset. To focus the analysis on leukemic blasts, we removed shared clusters containing cells that came from multiple patients (see associated code for details).

For differential expression analysis, leukemic blasts counts from a given patient sample were aggregated into pseudobulk profiles/samples (Squair et al., 2021). Differential expression analysis was performed on pseudobulk data using the likelihood ratio test (LRT) in DESeq2 v1.36.0. Genes with adjusted p-value < 0.05 were considered statistically significantly differentially expressed.

### Classifying B-ALL subtypes using explainable machine learning model

A neural network model was developed using TensorFlow (Keeton, 2016) in Python to classify B-cell Acute Lymphoblastic Leukemia (B-ALL) subtypes. We used 87 previously published single cell B-ALL samples collected at diagnosis from project SCPCP000008 (Iacobucci et al., 2025) in the Alex’s Lemonade Stand Single-cell Pediatric Cancer Atlas (Hawkins et al., 2025) for model training. After filtering out clusters of cells presumed to be non-leukemic blasts based on co-clustering across patients (see code), model input data consisted of a gene expression matrix with 751,904 cells and 22,261 genes. We split the data into training and testing using 80:20 split resulting in 601,523 cells for training and 150,381 cells for testing across 14 subtypes. These data were then normalized using MaxAbsScaler from the scikit-learn (Pedregosa et al., 2011) package to scale feature values appropriately while preserving sparsity in the expression data to accommodate datasets processed in two different batches (pilot and s2022).

We trained a neural network with a dense hidden layer with 500 neurons and ReLU activation, an output SoftMax layer with categorical cross-entropy loss using Adam optimizer with learning rate of 0.001. We used a batch size of 1000 with early stopping (patience of 10 epochs monitoring validation loss). While training, we further split the training dataset to training and validation with 80:20 split to monitor training performance using validation loss function. Model performance was evaluated using multiple metrics including F1 score, accuracy, precision, and recall on the test dataset.

We used SHAP (SHapley Additive exPlanations) (Lundberg & Lee, 2017), a popular feature importance package, to filter out non-important features reducing the number of genes (features) needed for subtype prediction. After extracting the top features, we trained a new model using the same architecture and evaluated the performance on test data. We deployed the model as a Streamlit app and as a CLI version for public use. We then tested the model against our single cell dataset from 25 patient samples with 79,216 cells. We made predictions for all the cells and implemented a majority voting approach for patient-level predictions, where the most frequently predicted class across all cells from a patient determined the final B-ALL subtype.

### Data and code availability

All raw data are available through the Childhood Cancer Data Sharing initiative at https://datacatalog.ccdi.cancer.gov/dataset/CCDI-phs002529. Processed single-cell data are available through the Single Cell Portal at https://singlecell.broadinstitute.org/single_cell/study/SCP3385/single-cell-profiling-of-pediatric-leukemia-patient-samples. Code for the analysis performed in this paper is available at https://github.com/ChildrensMercyResearchInstitute/ped_leukemia_RNA2025. Code for the single-cell classifier is available at https://github.com/ChildrensMercyResearchInstitute/cm-scbll.

## Results

### An institutional biorepository facilitates the study of transcriptional profiling across a diverse pediatric leukemia cohort

To identify key transcriptional characteristics of pediatric leukemias, we utilized the samples and associated clinical data available in the Children’s Mercy Cancer Biorepository (CM Tumor Bank). A total of 100 samples from 69 patients were profiled using bulk and/or single-cell RNA sequencing (Supplementary Table 1). Patients were selected to represent a broad set of leukemias and molecular subtypes, including a number of cases that had been profiled but could not be classified into a known genetic subtype. Bulk and single-cell RNAseq data were processed using standard open-source tools, including DESeq2 for bulk and Seurat for single-cell analysis. We then performed integrative analyses between the datasets to identify both common and subtype-specific transcriptional features of pediatric leukemia (Figure 1A).

**Figure 1.**
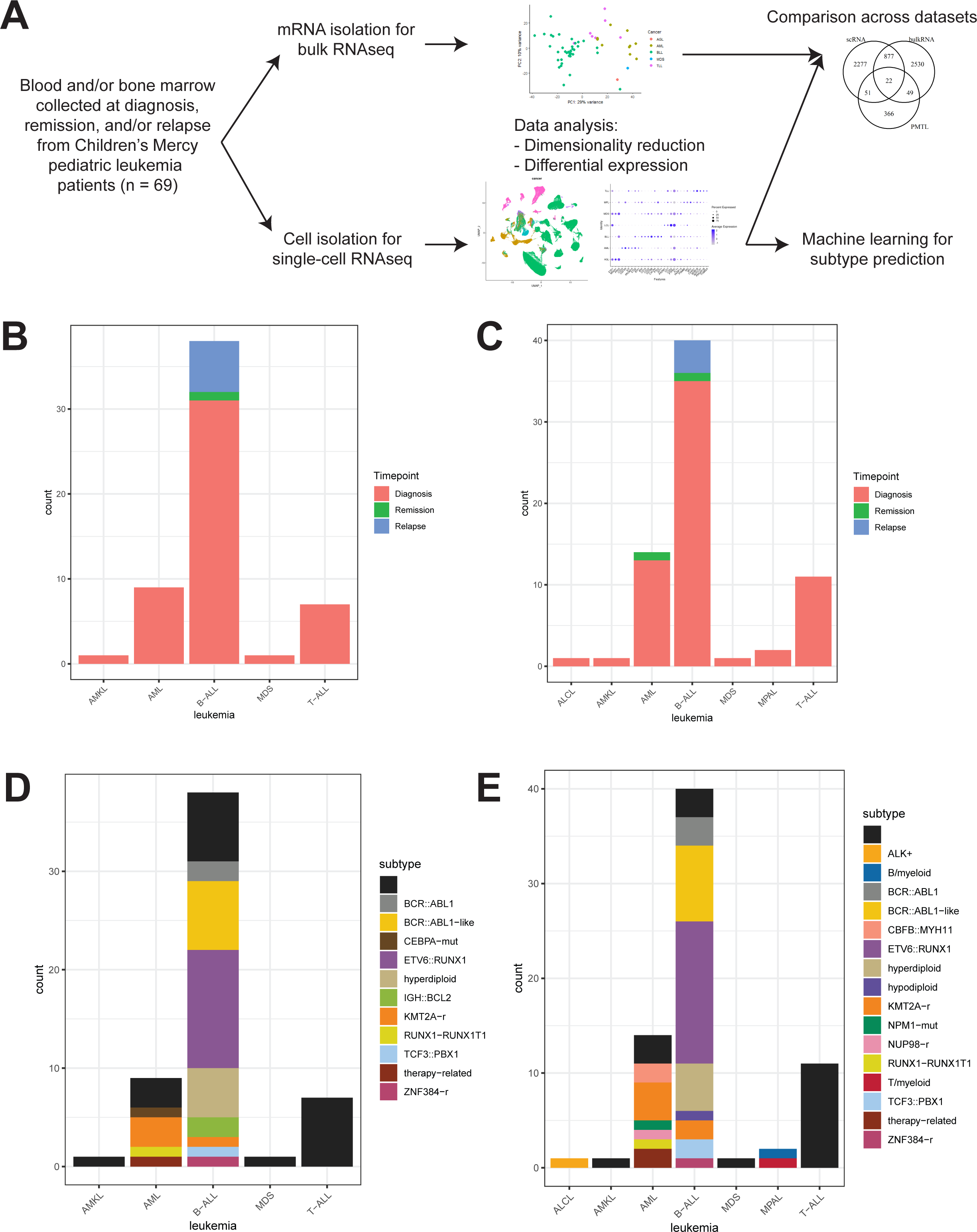
A) Schematic of sample collection and processing. B) Distribution of 56 pediatric leukemia patient bulk RNAseq profiles according to timepoint. Note that 6 patients have multiple samples, either across timepoints or tissue types (blood vs. bone marrow). C) Distribution of 70 pediatric leukemia patient single-cell RNAseq profiles according to timepoint. Note that 12 patients have multiple samples, either across timepoints or tissue types. D-E) Molecular subtype distribution across bulk (D) and single-cell (E) patient sample profiles. Full sample details can be found in Supplementary Table 1.

We generated bulk RNAseq data for a total of 56 blood or bone marrow samples and single-cell RNAseq data for a total of 70 blood or bone marrow samples from pediatric leukemia patients enrolled in the CM Tumor Bank. B-cell acute lymphoblastic leukemia (B-ALL) was the most frequently represented leukemia, followed by acute myeloid leukemia (AML) and T-cell acute lymphoblastic leukemia (T-ALL). Most of the samples in both bulk and single-cell datasets were collected at patient diagnosis (Figure 1B-C). For leukemias with defined molecular subtypes, we see similar distributions between bulk and single-cell datasets (Figure 1D-E).

### Bulk RNAseq reveals transcriptional similarities and distinct features across pediatric leukemias

We began by exploring the bulk RNAseq profiles of diverse leukemias. Transcriptional similarity across samples appears to be primarily driven by leukemia type (Figure 2A), highlighting the diagnostic and biological relevance of cancer-specific gene expression profiles. Tissue of origin (blood or bone marrow) did not appear to be a key discriminator in sample transcriptional identity (Figure 2B). In four cases where both sample types were collected from the same patient at the same clinical timepoint, expression profiles were highly similar, as indicated by the close proximity of paired samples (connected by lines in Figure 2B). This suggests that, within an individual, leukemia transcriptional identity remains consistent across sampling sites.

**Figure 2.**
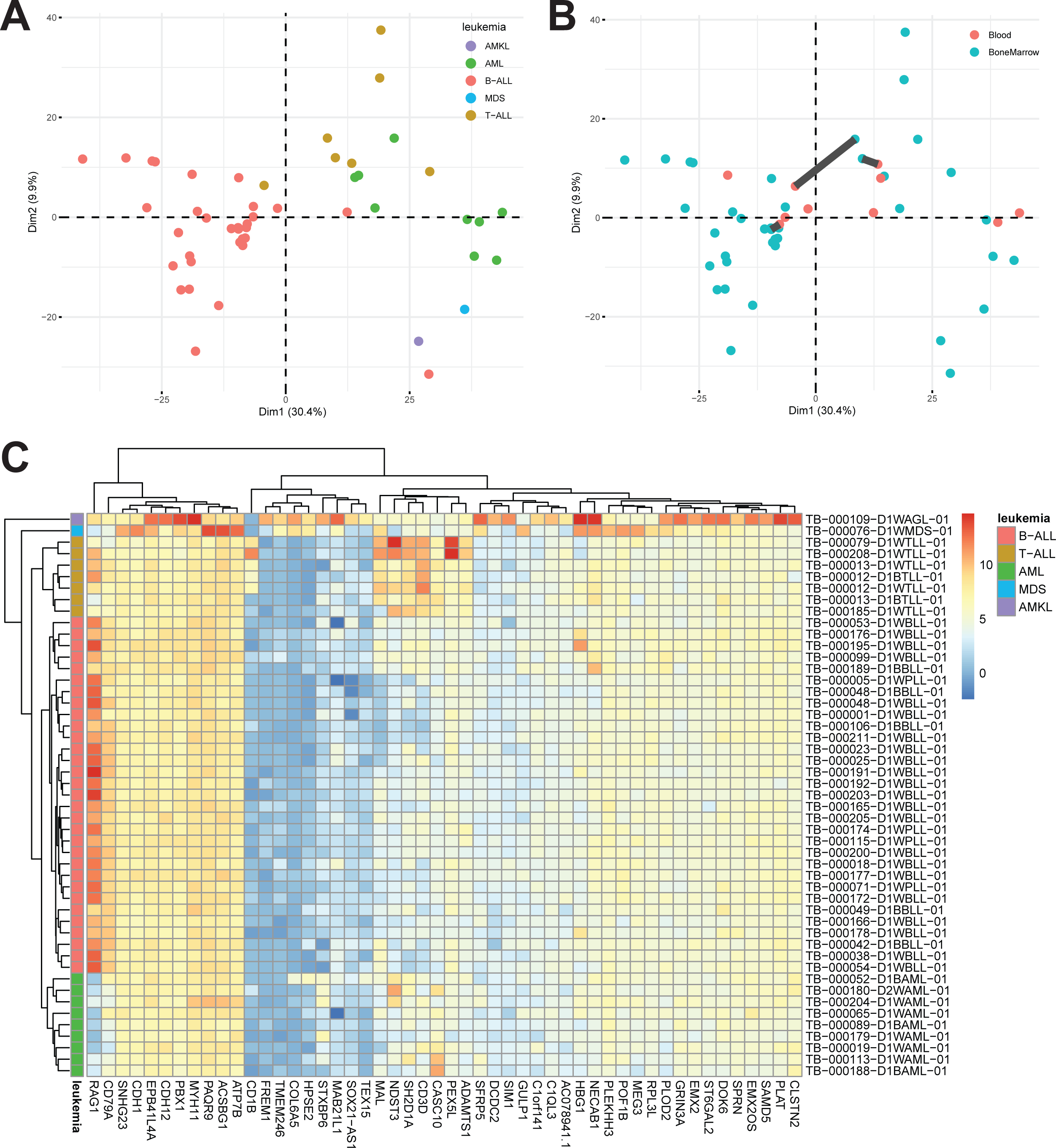
A) PCA plot depicting samples colored according to leukemia. B) PCA plot depicting tissue of origin. Patients with both a blood and bone marrow sample connected. C) Heatmap of expression (log normalized counts) of top 50 leukemia-specific genes.

To identify leukemia-specific signatures at diagnosis, we performed differential expression analysis across the represented leukemias, yielding 6,620 differentially expressed genes with p < 0.05 (Supplementary Table 2). Visualization of the top 50 genes (by adjusted p-value) shows strong leukemia-specific patterns of gene expression (Figure 2C). Additionally, hierarchical clustering of all samples collected at diagnosis mirrors transcriptional similarity patterns observed in PCA, with the single acute megakaryoblastic leukemia (AMKL) and myelodysplastic syndrome (MDS) samples, as well as one B-ALL sample, clustering with AML samples.

Focusing on a subset of genes to get a better sense of the heterogeneity of transcriptional findings, we looked at the intersection of genes differentially expressed across leukemias with the FDA’s Pediatric Molecular Target Lists (PMTL, MTP PMTL Documentation | Molecular Targets Platform, n.d.). A total of 160 genes with known roles in pediatric cancers showed a leukemia-specific pattern of expression in our data, including *FOXO1*, *CD7*, *ETS1*, and *KIT* (Supplementary Figure 1). Despite a high degree of variability within each leukemia, the overall distributions of gene expression showed leukemia-specific trends, supporting the power of this analysis to identify novel transcriptional signatures of interest.

### Single-cell RNAseq reveals heterogeneity across pediatric leukemias

Given the heterogeneous nature of cancer, we set out to explore the transcriptional profiles of these patient samples at higher resolution using single-cell RNA sequencing (scRNAseq). Consistent with prior single-cell studies of leukemias (Candelli et al., 2022; Caron et al., 2020; Ferrall-Fairbanks et al., 2019; Iacobucci et al., 2025), we observed that cell clustering is primarily driven by patient identity, with proximity in UMAP space also driven by cancer (Figure 3A-B). Based on ENCODE data, we used the SingleR package to assign each cell an identity based on transcriptional similarity to healthy cells (Figure 3C).

**Figure 3.**
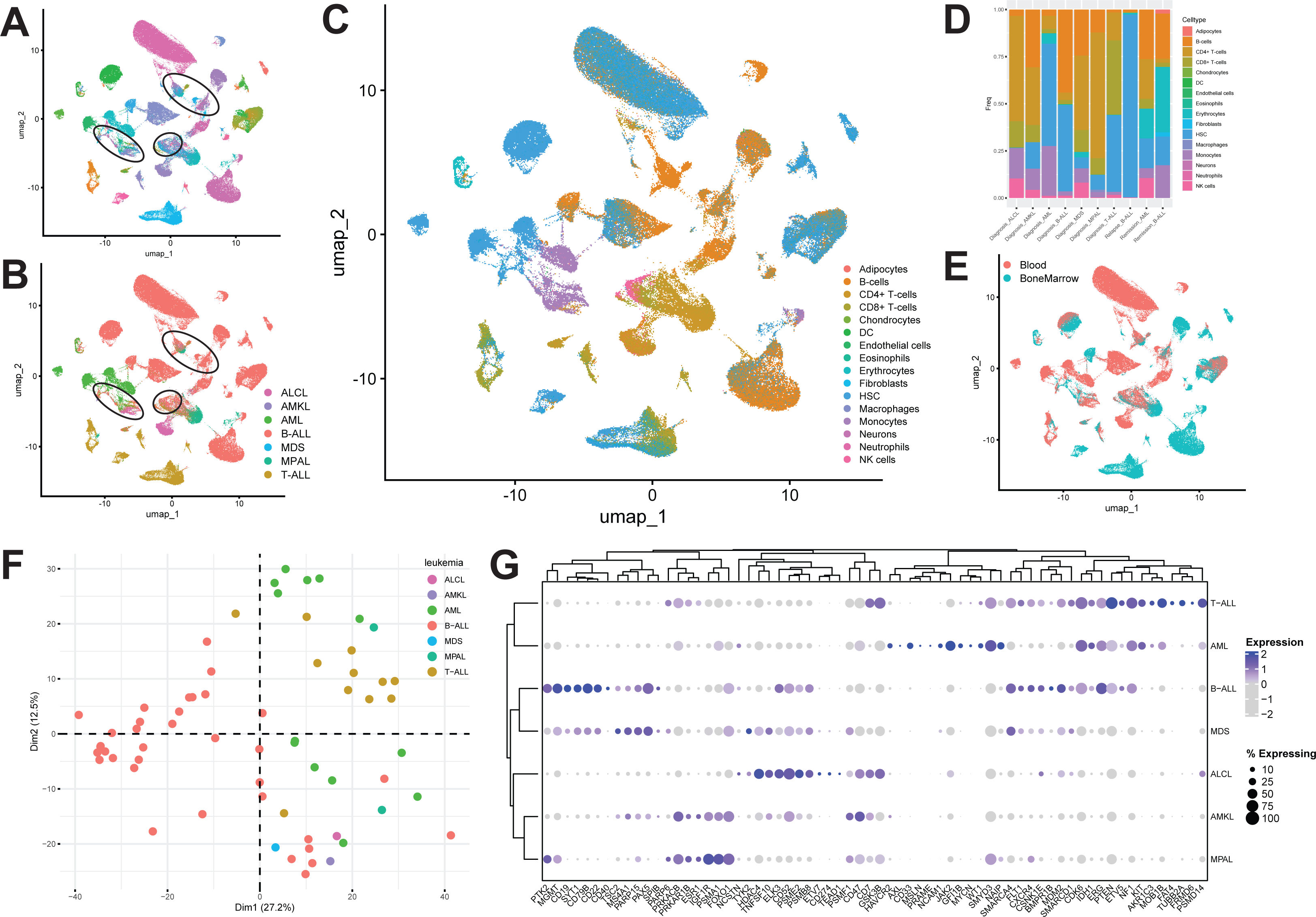
A) UMAP with cells colored according to patient of origin. Circled clusters correspond to mature (non-leukemic) cells. B) Cells colored according to patient leukemia diagnosis. C) Cell type as assigned by the SingleR package using ENCODE dataset. D) Proportion of cells for each leukemia at each timepoint most closely resembling each healthy cell type. E) Cells colored according to tissue of origin. F) PCA plot of pseudobulk leukemic cells colored according to leukemia. G) DotPlot depicting expression of genes differentially expressed across B-ALL subtypes in both bulk and single-cell pseudobulk datasets, which overlap with the PMTL.

We observed that mature cells tended to cluster together regardless of the patient of origin (circled clusters in Figure 3A-B), while leukemic cells form patient-specific clusters. Among the leukemic cell clusters, we saw high similarity to cells of origin for each leukemia (Figure 3C-D). For example, leukemic clusters from B-ALL patients contain high proportions of B-cell-like cells, while T-ALL cases predominantly show high proportions of T-cell-like cells. Across all leukemic clusters, we saw hematopoietic stem cell (HSC)-like cells, although the proportion of these cells varies across patient samples. Consistent with our observations from the bulk RNAseq (Figure 2A), we saw a tendency for patient clusters to occupy adjacent regions in UMAP space based on the cancer type (Figure 3B). Similarly, the tissue type did not inherently inform clustering, but samples of different tissues from the same patient formed adjacent clusters (Figure 3E).

The resolution of scRNAseq allowed us to focus our analysis specifically on the cancer cells of a sample, eliminating artifacts due to varying blast percentages across samples and the presence of non-leukemic cells. Because mature cells clustered together in UMAP space regardless of the individual sample or cancer of origin, we manually selected such clusters and removed them from our dataset to leave a near-pure population of cancer cells (Supplementary Figure 2; see code). The use of multiplexing also allowed for better removal of ‘doublets’ given the greater likelihood that droplets containing two cells would be of different genotypes. For consistency with our processing of bulk RNAseq data and because of improved biological significance (Squair et al., 2021), we aggregated data across all cancer cells for a given sample and used the subsequent ‘pseudobulk’ samples for downstream analysis.

We observed transcriptional similarity across these pseudobulk samples (Figure 3F) that support the trends in our bulk RNAseq data (Figure 2A) and single-cell dimensionality reduction (Figure 3B), with patient samples showing separation according to cancer diagnosis. As with our bulk RNAseq data, we set out to identify leukemia-specific transcriptional signatures at diagnosis. Following the same differential expression testing methodology using our pseudobulk samples as for our bulk RNAseq analysis, we identified 4,795 genes showing a leukemia-specific pattern of expression with p < 0.05 (Supplementary Table 3).

Of these, 1,958 genes overlap with those identified from bulk RNAseq data. These genes are highly enriched for pathways expected to be involved in leukemia development and progression, including Hematopoietic cell lineage (KEGG:04640), Transcriptional misregulation in cancer (KEGG:05202), and cell differentiation (GO:0030154), supporting their roles in cancer-specific processes (Supplementary Table 4). Overlap of these genes with the PMTL and further filtering identified a set of genes with leukemia-specific expression patterns across leukemic cells (Figure 3G).

### Known genetic drivers result in subtype-specific transcriptional patterns in B-ALL

Both our bulk and single-cell RNAseq datasets contain leukemic samples from patients with a diverse set of molecular subtypes, as well as patients without clear driver mutations (Figure 1 D-E). We set out to explore transcriptional differences across molecular subtypes at diagnosis. To avoid confounding factors of leukemia-specific effects, we focused on samples collected at diagnosis from patients with B-ALL (bulk n = 31, single-cell n = 35), as it was the cancer with the best representation in our dataset.

Across both bulk and single-cell pseudobulk data, we observed that samples with a particular subtype tend to show transcriptional similarities (Figure 4A-B), consistent with the shared molecular drivers of disease in these patients. Strikingly, patients with *ETV6::RUNX1* show the strongest subtype-specific clustering in both datasets. Patients lacking clear driver mutations show up throughout PCA and correlation space, showing varying degrees of transcriptional similarity to patients with known drivers.

**Figure 4.**
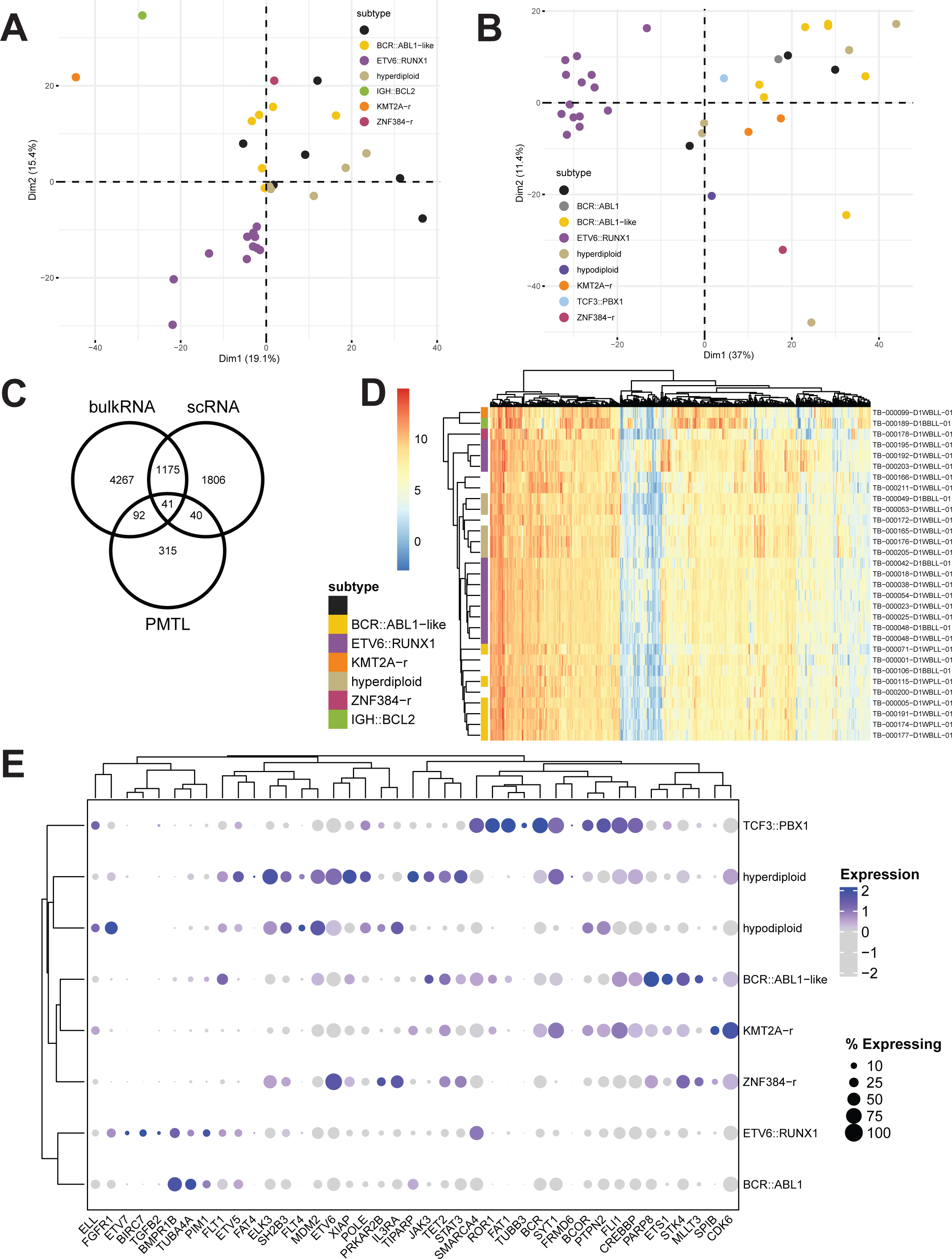
A-B) PCA plots demonstrating transcriptional similarity of B-ALL subtypes in bulk (A) and single-cell pseudobulk (B) datasets. C) Venn diagram depicting overlap of genes differentially expressed across B-ALL subtypes in scRNAseq data, bulk RNAseq data, and found on the FDA’s PMTL list. D) Expression heatmap (log normalized counts) of top 500 genes differentially expressed in both bulk and scRNAseq data. E) DotPlots of gene expression demonstrating subtype-specific signatures – 41 genes differentially expressed in bulk, scRNA data, and appearing on PMTL list.

To further explore the molecular underpinnings of subtype-specific transcriptomes, we performed differential expression analysis across all patients with known genetic subtypes. Bulk RNAseq data identified 5,575 genes differentially expressed with adjusted p-value < 0.05, while pseudobulk single-cell data identified 3,062 genes (Supplementary Tables 5-6). 1,216 genes were differentially expressed across subtypes in both bulk and pseudobulk datasets. It is important to note that perfect overlap is not expected, with some subtypes (hypodiploid and *TCF3::PBX1*) represented only in the pseudobulk dataset.

Of these genes, 41 further overlapped with the FDA’s PMTL list, including *JAK3*, *ETS1*, *PRKAR2B*, and other genes known to play key roles in specific subtypes of leukemias (Figure 4C). Both bulk (Figure 4D) and single-cell (Figure 4E) data showed subtype-specific expression of many of the genes identified through this analysis. This facilitates the establishment of subtype-specific gene expression signatures and enables the development of mechanistic hypotheses based on expression of known and novel marker genes.

### Single-cell transcriptomes facilitate the characterization of patient samples with unknown cancer drivers

The utility of our rich transcriptomic dataset extends beyond the characterization of pediatric leukemias with well-defined molecular drivers. To highlight its potential in analyzing patient samples lacking clear genomic alterations, we developed a neural network model trained on single-cell transcriptomic data, which we then tested on this dataset. As detailed in the Methods and illustrated in Figure 5A, the model was trained using 87 pediatric B-ALL samples at diagnosis from a previously published study (Iacobucci et al., 2025). Given the sparsity of single cell data, we used SHAP to identify the top 10,000 most informative transcriptomic features for accurately classifying 14 distinct B-ALL subtypes.

**Figure 5.**
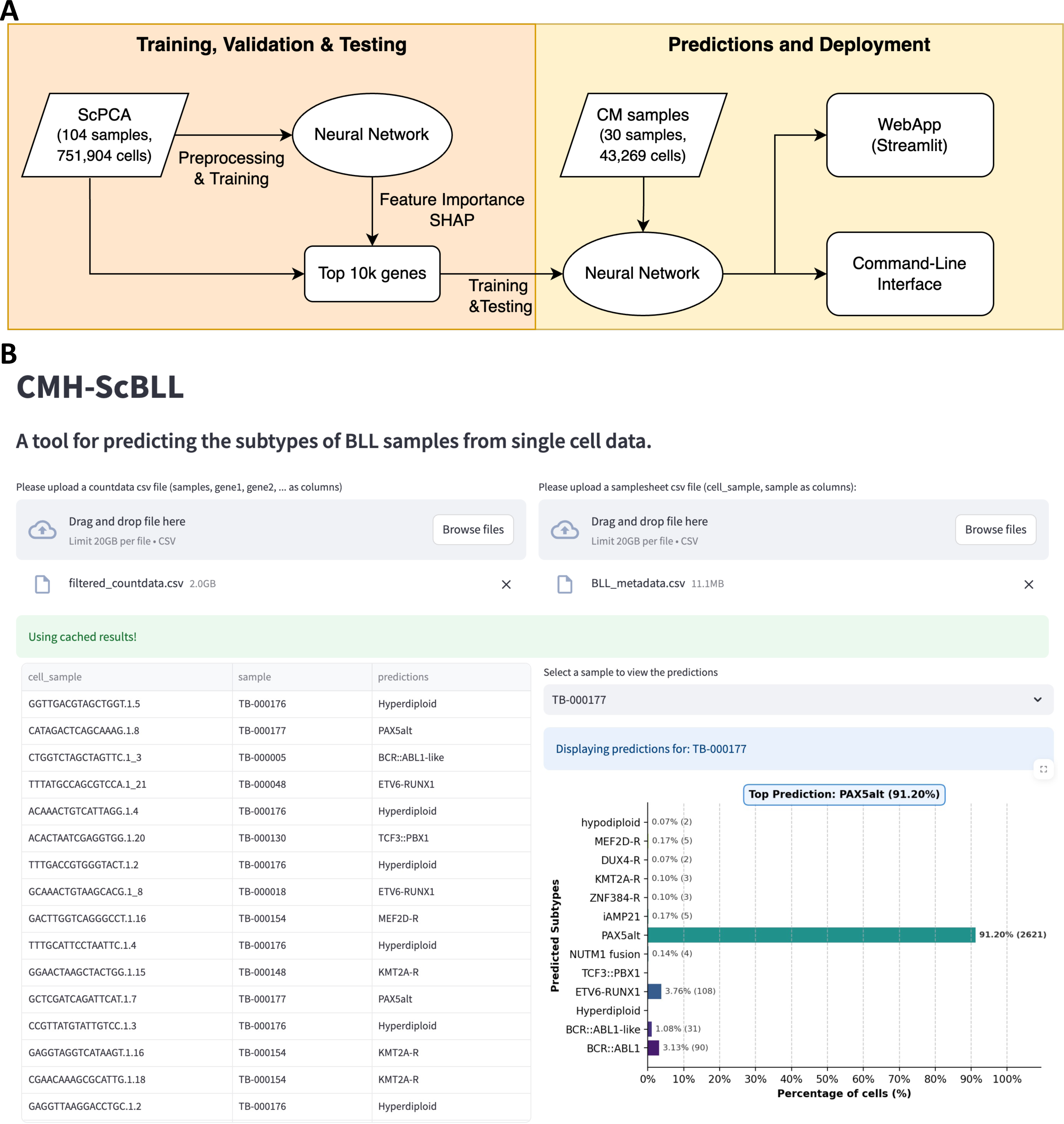
A) Schematic depicting explainable machine learning model training and testing using training data from Iacobucci et al. and testing on CM samples. B) Streamlit web app for uploading single cell count data and metadata to generate predictions.

We achieved high performance on the held-out test set derived from the Iacobucci et al. dataset, with an F1-score of 0.99 (Table 1). When applied to the CM cohort, our model achieved an F1-score of 0.81 at the cell level and 0.84 at the patient level. Detailed prediction metrics for each subtype are provided in Supplementary Table 7. Subtype predictions for CM samples at both the cell and patient levels are presented in Supplementary Table 8, including potential subtype resolution in previously unclassified cases.

Among the 25 patients in the CM cohort, two were unclassified, and our model was able to assign subtypes to two patients who did not have a confirmed diagnosis at the time of initial analysis. Specifically, patients TB-000001 and TB-000166 were initially categorized as NOS (not otherwise specified). Our model predicted TB-000001 as belonging to the DUX4 subtype. This prediction was supported by testing performed after initial classification, indicating IGH FISH positivity and the presence of a *DUX4* translocation. For TB-000166, no fusions were detected, and FISH results were negative. The presence of an *IKZF1* deletion in clinical testing often correlates with a Ph-like classification. However, our model predicted it as 56% (209/376 cells) Hypodiploid and 35% (130/376 cells) iAMP21 subtypes. This lack of concordance may be due to the low number of cells sequenced from this sample.

Two additional classification differences were observed in patients TB-000115 and TB-000177, both previously diagnosed as Ph-like. The model predicted both as PAX5alt with 53% (236/443) and 91% (2621/2874) of cells, respectively. Interestingly, both cases showed some subtyping ambiguity. Despite extensive clinical testing, no common Ph-like driver fusion was detected, though the samples showed partial CRLF2+ in flow cytometry and was positive for Ph-like gene expression and high *CRLF2* expression in the Tricore Low Density Array assay. For TB-000115, a SNP-based microarray to assess for copy number changes at the DNA level also showed a heterozygous loss of *PAX5*, which may explain the classification. Similarly, TB-000177 showed both *P2RY8::CRLF2* fusion, a common Ph-like drivers, as well as a gain of *PAX5* (with a Tier II SNV in the amplified region), which likely affected model performance. Interestingly, similar cases with both *PAX5* amplification and *P2RY8::CRLF2* fusion have been observed (Schwab et al., 2017). Using the B-ALL gene expression classifier ALLCatchR (Beder et al., 2022), this case was also classified as PAX5alt (though with a score of 0.55 on a 0-1 scale, and a categorization of “candidate” as opposed to “high-confidence”). It appears that both of these cases may show transcriptional programs with both Ph-like and PAX5alt characteristics, demonstrating the value of multiple assays for subtyping.

Overall, this modeling approach quantifies the transcriptional similarity of leukemia samples with unknown drivers to established B-ALL subtypes, offering insights into the molecular basis of individual cases. The model is accessible via a web interface (Figure 5B), allowing users to generate subtype probability scores at both the cell and patient levels across 14 B-ALL subtypes. Code is publicly available at https://github.com/ChildrensMercyResearchInstitute/cm-scbll. This predictive tool underscores the utility of high-resolution transcriptomic data for investigating the etiology and progression of pediatric leukemias.

## Discussion

In recent years, cancer profiling has increasingly moved beyond the genome to integrate additional data gained from transcriptomic analysis in both research and clinical applications. While data from pediatric cohorts has always been in shorter supply than adults, novel efforts to collect and harmonize pediatric transcriptional data are beginning to flourish, including the NCI’s Childhood Cancer Data Initiative (Flores-Toro et al., 2023; Jagu et al., 2024), Gabriella Miller Kids First (Hudson et al., 2023), Alex’s Lemonade Stand Foundation’s Single-cell Pediatric Cancer Atlas (Hawkins et al., 2025), and the Treehouse Initiative (Bjork et al., 2019). A current downside of these repositories is that they are often populated with individual datasets that focus on a specific cancer type, and make cross-dataset, and thus cross-disease, comparisons challenging due to batch effects.

Here, we present a large dataset of bulk and single-cell pediatric leukemia transcriptomes. The data span a total of eight leukemias across three treatment timepoints, and facilitate integrated, comparative analyses of these cancers (Figure 1). To date, this is one of the largest pediatric leukemia cohorts to be published, which facilitates an unprecedented level of comparison across both bulk and single-cell data, supplementing existing resources (Iacobucci et al., 2025; Mumme et al., 2022; Pölönen et al., 2024; Sanders et al., 2019; Tan et al., 2023; Vaske et al., 2019).

We performed comprehensive transcriptional comparisons across the leukemias represented in the bulk and single-cell datasets, identifying key genes more highly expressed in each leukemia (Figure 2-3). We were able to recapitulate prior findings including overexpression of genes of interest relative to other leukemias – *FOXO1* in B-ALL (Bhansali et al., 2021), *MSLN* in AML (Le et al., 2021), and *ETS1* in T-ALL (McCarter et al., 2020). With a high degree of overlap between bulk and single-cell differential expression analysis, novel leukemia-specific genes of interest can be investigated to explore potential molecular mechanisms involved in driving each disease.

To demonstrate the utility of these data toward clinical applications, we performed further analysis of pediatric B-ALL transcriptomes to discriminate between well-defined molecular subtypes. We identified genes of interest expressed in subtype-specific patterns across both bulk and single-cell RNAseq datasets (Figure 4). A number of the genes identified through this analysis are targetable with existing therapies, including *STAT3* (Zhu et al., 2023), *FLT3* (Brown et al., 2021; Suematsu et al., 2023), and *BCL2* (Pariury et al., 2023). The variability of expression of these genes across B-ALL subtypes is not surprising but has significant implications for treatment selection for pediatric B-ALL patients. These findings present an opportunity for mechanistic follow-up to establish subtype-specific gene expression signatures and corresponding treatment options.

Finally, we leveraged existing data (Iacobucci et al., 2025) and our scRNAseq B-ALL dataset to train an explainable machine learning model for predicting the similarity of individual cells to known B-ALL subtypes (Figure 5). Although classifiers have been published that use bulk RNAseq data (Beder et al., 2022; Gu et al., 2023; Krali et al., 2023; L. Li et al., 2025; Schmidt et al., 2022), our use of scRNAseq data enables deeper exploration of the heterogeneity of these cancers. Significantly, our work also supports the value of integrating RNA-based profiling (both bulk and single-cell) with DNA-based clinical assays to improve our understanding of the molecular landscapes driving individual patients’ leukemias.

In summary, here we present bulk and single-cell RNAseq data from a large cohort of pediatric leukemia patient samples to characterize distinct leukemias and molecular subtypes of B-ALL. We demonstrate that our data recapitulate known facets of pediatric leukemia biology and present the opportunity to develop novel mechanistic hypotheses based on genes of interest identified through this analysis. Additionally, we show concordance between our transcriptional profiling efforts and clinically significant genomic findings, suggesting avenues toward future application of transcriptomics in a clinical setting. The data and approaches presented here represent a valuable addition to the field of pediatric leukemia, with applications in both research and the clinic.

## Supplementary Data

**Supplementary Figure 1.** Leukemia-specific expression of representative genes of interest in bulk RNAseq data.

**Supplementary Figure 2.** Visualizations of clusters removed to focus downstream analysis on leukemic blasts – left is full Seurat object, right is with “healthy/mature” cells removed. Colored according to A) patient of origin, B) leukemia, and C) closest healthy cell type assignment using ENCODE dataset.

**Supplementary Table 1.** Sample metadata including diagnosis, sex, leukemia (and subtype, if known), and sequencing batch for bulk and single-cell RNAseq.

**Supplementary Table 2.** Differentially expressed genes for bulk RNAseq across leukemias.

**Supplementary Table 3.** Differentially expressed genes for single-cell RNAseq (using pseudobulk approach) across leukemias.

**Supplementary Table 4.** Pathway enrichment analysis results of all genes differentially expressed across leukemias in both bulk and single-cell RNAseq.

**Supplementary Table 5.** Differentially expressed genes for bulk RNAseq across B-ALL subtypes.

**Supplementary Table 6.** Differentially expressed genes for single-cell RNAseq (using pseudobulk approach) across B-ALL subtypes.

**Supplementary Table 7.** Performance metrics of Sc-BLL model on test data (split from ScPCA) and CM datasets across 14 B-ALL subtypes.

**Supplementary Table 8.** Subtype prediction level of Sc-BLL model at cell and sample level for all CM samples.

## Acknowledgements

We would like to acknowledge the Children’s Mercy Research Institute Biorepository (CRIB) for assistance with sample collection and processing. This work was supported by an MCA Partners Advisory Board grant from Children’s Mercy Hospital (CMH) and The University of Kansas Cancer Center (KUCC).

## Funding

This work is funded by the contributions from these below funds:

Elizabeth and Monte McDowell

Big Slick

Masonic Cancer Alliance

Black & Veatch Foundation

Curing Kids Cancer

Braden’s Hope for Childhood Cancer

Alex’s Lemonade Stand

Children’s Mercy Cancer Center Auxiliary

