## Supplementary figures and images for "A comprehensive bulk and single-cell transcriptional atlas of pediatric leukemias"

### Supplementary Figure 1

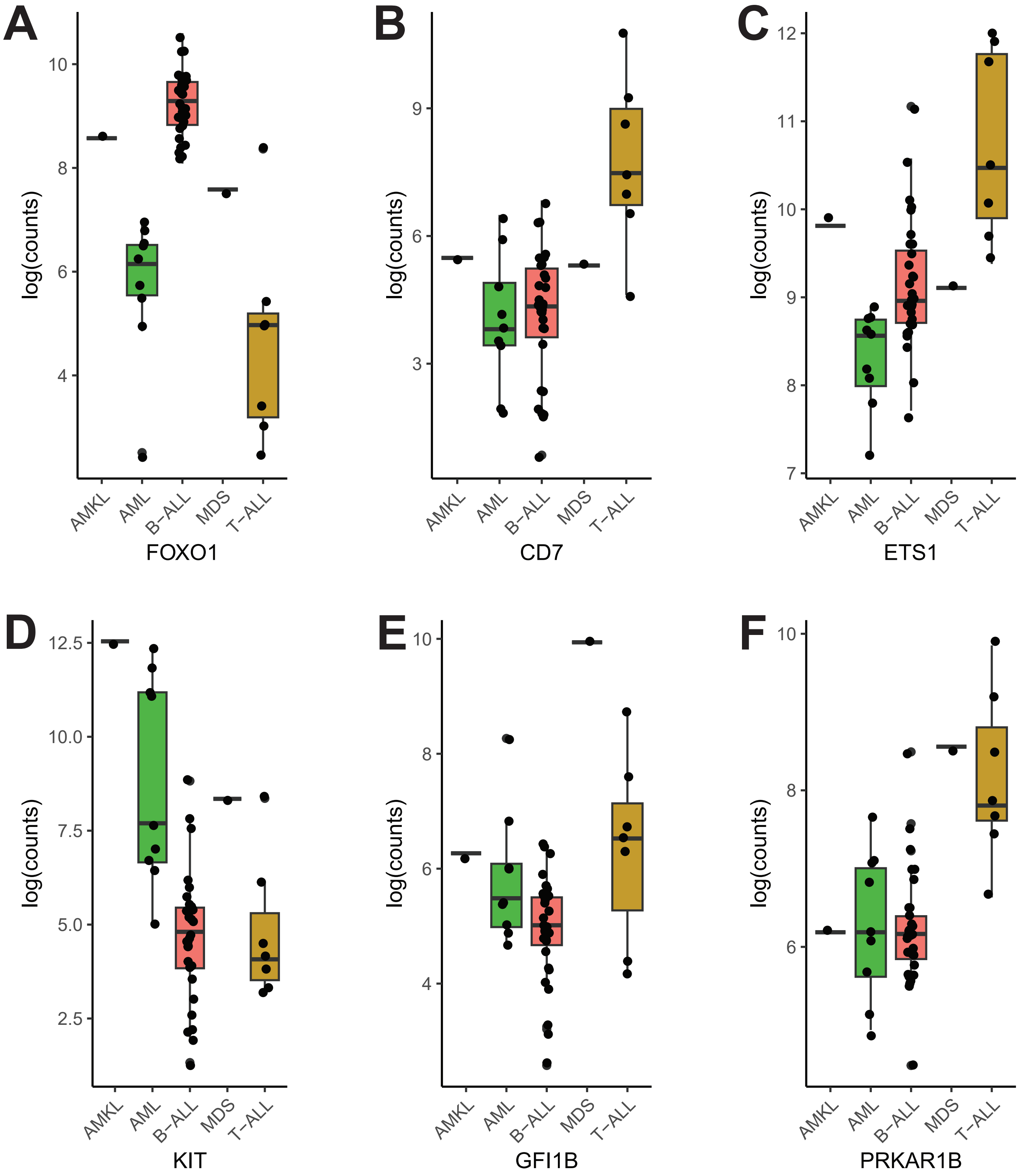

### Supplementary Figure 2

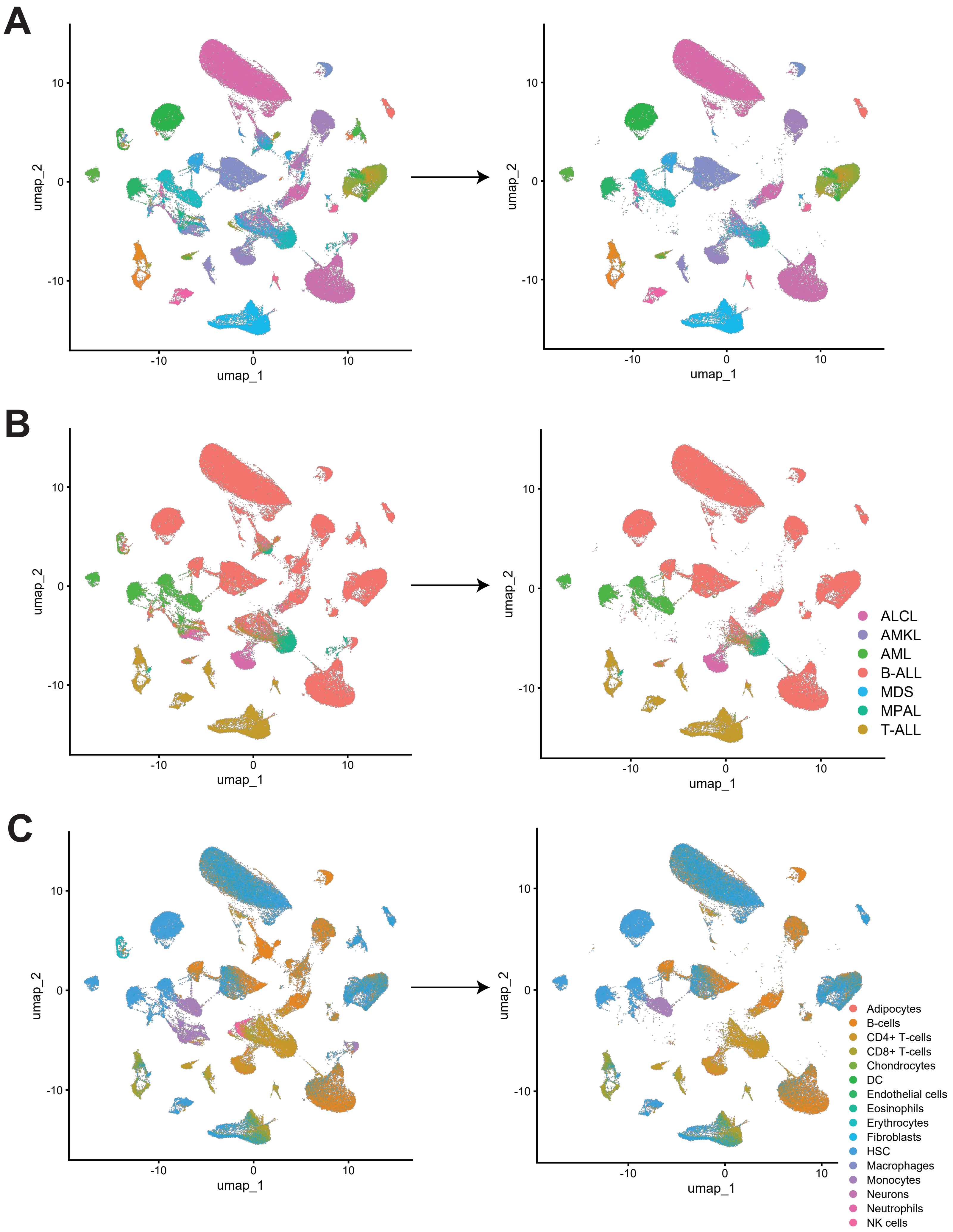
